# Role of charge in enhanced nuclear transport and retention of graphene quantum dots

**DOI:** 10.1101/2023.09.27.559706

**Authors:** Gorav Gorav, Vrushali Khedekar, P. Nandakumar, Geetha K. Varier

## Abstract

The nuclear pore complexes on the nuclear membrane function as the sole gateway of molecular communication between the nucleus and the cytoplasm regulating the transport of molecules, including nucleic acids and proteins. The present study seeks to undertake a comprehensive investigation of the kinetics of transport of negatively charged graphene quantum dots through nuclear membranes and quantify their nuclear transport characteristics and translocation rates. Experiments are carried out in permeabilized HeLa cells using time-lapse confocal fluorescence microscopy. Introducing negative charge onto biomolecular probes leads to electrostatic interaction with the nuclear pore complexes resulting in significant changes in their nuclear translocation rates. We find that the negatively charged graphene quantum dots are transported to the nuclei at a fast rate and two distinct transport pathways are involved in the translocation. Furthermore, complementary experiments on the nuclear import and export of these graphene quantum dots confirm the bidirectionality of transport with similar translocation rates. Our studies also show that negatively charged graphene quantum dots exhibit good retention properties revealing their potential as excellent drug carriers.

**Statement of significance:** The nuclear pore complexes control the bidirectional transport of biomolecules between the nucleus and cytoplasm. The noteworthy behaviors exhibited by negatively charged graphene quantum dots with respect to the nuclear uptake show their potential utility not only as drug carriers but also as facilitators for the retention of drugs within the nucleus. The fast import of carriers helps to achieve faster drug delivery, and the retention ensures the passing of the drug to daughter nuclei.

**Graphical abstract:** 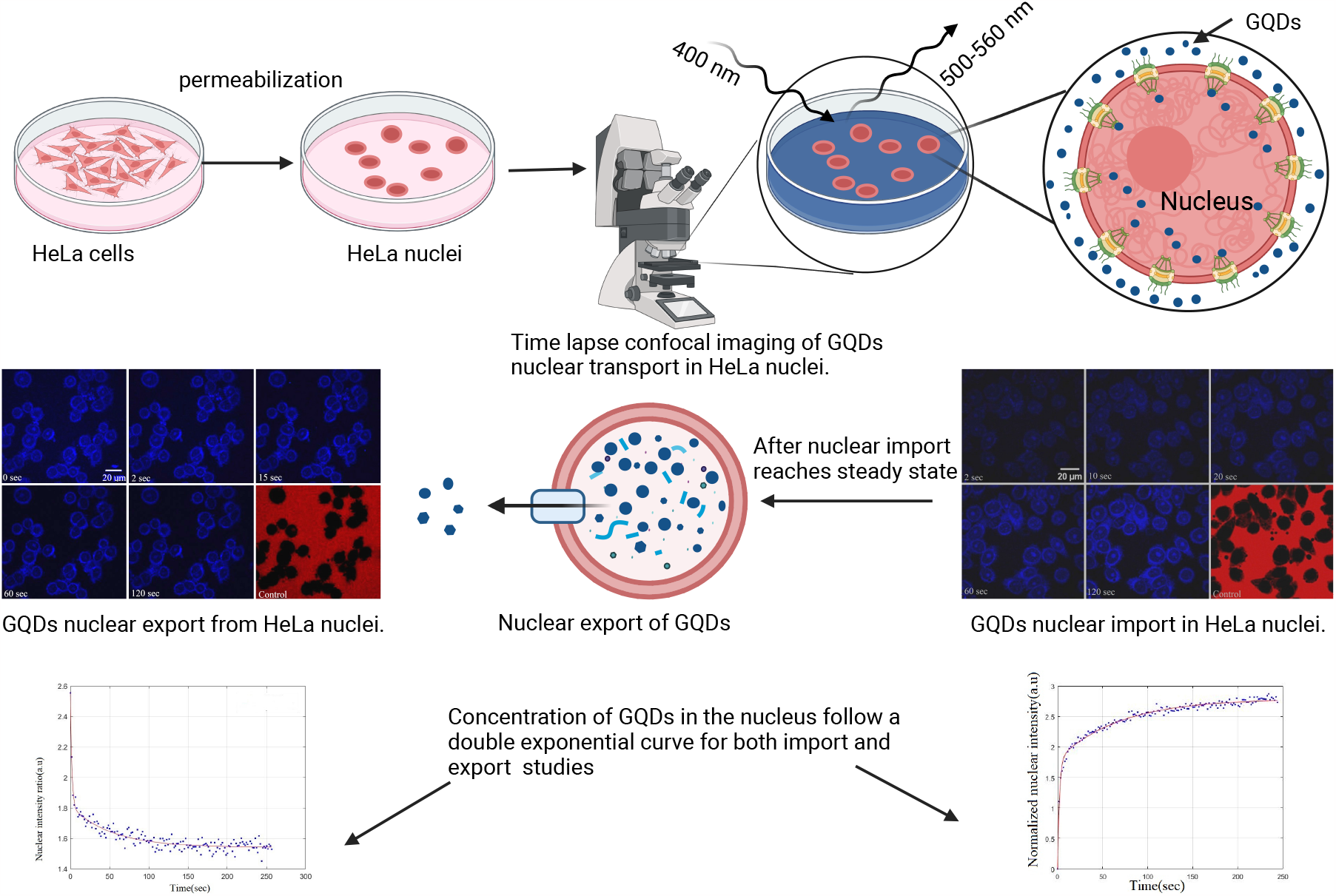

## 1 Introduction

The nucleus is a specialized organelle of eukaryotic cells that harbours the cell’s genetic material in the form of chromatin, a complex of DNA and proteins. The nucleus plays crucial roles in regulating gene expression, DNA replication, DNA repair, and cell cycle progression. The nuclear envelope is a dynamic structure that separates the nucleus from the cytoplasm and regulates the nuclear import and export of molecules via the nuclear pore complexes (NPCs). NPCs are complex protein assemblies of size ≈ 125 MDa on the nuclear envelope [1, 2, 3, 4, 5]. The molecular transport between the nucleus and cytoplasm is mainly classified based on the size of the transporting molecule. The molecules having a molecular weight of less than 40 kDa or a diameter of 9 nm are transported to the nucleus through passive diffusion, whereas bigger molecules pass via an active transport mechanism [5, 6, 7, 8, 9]. There are around 3,000 – 5,000 NPCs in a proliferating human cell [10, 11]. The translocation capacity of NPCs is enormous, around 100 MDa/s. The active transport of cargo in and out of the nucleus is mediated by specific signals such as nuclear localization signal (NLS) and nuclear export signal (NES). Karyopherins (importins and exportins) possess binding sites for NLS and NES, and the complex they form with cargo translocates through NPCs by interacting with phenylalanine-glycine (FG) Nucleoporins (Nups) [12, 13, 5].

Nuclear translocation is an essential phenomenon in every living cell. To understand this complex process, we should know the interior structure of the NPCs. The NPCs have eight-fold symmetry on both cytoplasmic and nuclear sides and comprise approximately 30 different types of Nups. The human cell NPCs mainly consist of Nups 214, 88, and 62 towards the cytoplasmic side, Nups 98, 62, 58, and 54 on the central ring, and Nups 153 and 50 on the nuclear side of the nuclear membrane [14]. These Nups contain repeats of phenylalanine and glycine amino acids connected through hydrophilic linkers [1, 15, 2]. The structure of FG Nups can either be an extended coil, collapsed coil, or both, depending on the charge content of amino acids present in the Nups. One-third of FG Nups are intrinsically disordered proteins (IDPs), forming selective hydrogel mesh along the central axis and dynamic in nature [16]. During the process of nucleocytoplasmic transport, the cargo molecules restrict the flexibility of FG-IDPs leading to a decrease in entropy. Small molecules do not affect much of IDP’s entropy during diffusion through NPCs, but large molecules restrict the IDP’s movements and decrease the entropy of FG-IDPs. The hydrophobic interaction between cargo complex (cargo+NLS+ karyopherins) and IDPs can decrease the enthalpy of the system (FG Nups + cargo complex) to compensate with entropy decrease and is termed as enthalpy - entropy compensation [17, 18]. Small biomolecules can diffuse through the gaps between IDP structures. On the contrary, hydrophobic FG IDPs act as a barrier for large molecules (size more than 40kDa) [16, 9, 17]. Earlier, the only interaction known to help in overcoming the barrier was the hydrophobic interaction between the cargo complex and the FG Nups. Now several research groups have documented that electrostatic interaction between negatively charged cargo and positively charged FG Nups also contribute to nuclear transport [19, 20, 21, 22].

Studies on biomolecular transport through nuclear membranes have been an important technique for probing the nuclear pore complexes. Inert dextran molecules, available in different sizes, is one of the model systems widely employed to probe the mechanism of transport through nuclear membranes. Dextran molecules appropriately labeled with fluorescent dyes such as fluorescein isothiocyanate (FITC) or tetramethyl rhodamine isothiocyanate (TRITC) are used in these experiments to examine their nuclear uptake using confocal fluorescence microscopy. These studies have revealed valuable information on nuclear transport characteristics, including the size of the nuclear pores and the permeability properties of the nuclear membrane. The molecular weight of the transporting molecules significantly affects their nuclear uptake. [8, 6]. It is found that the nuclear accumulation and retention time of dextran inside the nucleus is linearly dependent on the molecular weight. [23]. Several studies carried out in tumor-bearing mice have confirmed that the electric charge significantly impacts the plasma and tissue disposition of dextrans. The positively charged dextran accumulates in tissues from plasma at a higher rate, whereas negatively charged dextran moves slowly from plasma to tissue [24, 25]. It is also observed that the rate of nuclear transport of dextran is affected by the excluded volume effect caused by molecular crowding in the nucleus. The human cell nuclei being 30-40 % filled with chromatins [26], the dextran concentration makes the nuclei more crowded and results in steric repulsion. It is noted that FITC dextran is excluded from the chromatin region and nucleoli. Molecular crowding depends on the interaction between the cargo and the environment. The cargo having weak interaction with the environment can overcome crowding behavior and steric repulsion [27].

In this context, nanoparticles, and quantum dots have grabbed a lot of research attention in recent years as potential biomolecular probes for nuclear transport studies. Nanoparticles are effective in targeted drug delivery and also reduce the side effects of chemotherapy [28, 29, 30, 31]. Quantum dots or nanoparticles can be used as a probe in different microscopic techniques such as two-photon microscopy, confocal microscopy, and fluorescence resonance energy transfer microscopy [32, 33, 29, 31, 34]. Due to the significant surface-to-volume ratio of nanoparticles, it is easier to attach peptides that help in cellular, nuclear, and endosomal membrane escape [28, 29, 35]. Studies show that the nanoparticle’s cellular and nuclear uptake depends on its elasticity, size, and surface properties [28, 29, 36, 30]. Gold nanoparticles were widely used in these research studies; now, graphene quantum dots (GQDs) have emerged as another potential probe of interest due to their low toxicity, biocompatibility, easy surface modification, the high reactivity of the edge, unique sp2 hybridized crystal structure, and oxygen-rich functional group. They are also useful for microscopic imaging due to their autofluorescence properties, high quantum yield, photo-stability, and reduced photo-bleaching [32].

In the present work, the kinetics of nuclear import and export of GQDs are studied using confocal fluorescence microscopy. Experiments are carried out in digitonin-permeabilized cell lines, one of the well-established systems for studying transport under transient conditions. Digitonin ruptures only the cell membrane during permeabilization and keeps the nuclear membrane intact, thus enabling us to study the transport of molecules directly through the nuclear membrane. GQDs, used as potential cargo molecules in our studies, contain a carboxyl group and are negatively charged. To the best of our knowledge, though different groups have observed the cellular and nuclear uptake of GQDs, the kinetics of transport of these quantum dots are not well studied, and their translocation rates are not yet quantified. The maximum amount of GQDs that can be accommodated in the nucleus is not known so far. Our studies reveal the rate constants of transport of GQDs through the HeLa cell nuclear membrane and throw light on many interesting facets of the nuclear import and export of GQDs. %The images shown are representative of one experiment from a repetition of many experiments.

Studies are conducted in HeLa cell nuclei in two different ways. In one set of experiments, an import mixture containing RRL (Rabbit reticulocyte lysate) along with cargo is used, while in another set of experiments, the import mixture that does not contain RRL is used. RRL, containing importin protein and transport factors, is used to mimic the cytoplasmic contents of the cell. The GQDs used in nuclear transport studies are negatively charged with a zeta potential value of -32.4 mV. The size of GQDs characterized using transmission electron microscopy and scanning electron microscopy was found to be approximately 6 nm and 10 nm, respectively. The hydrodynamic size observed through the particle size analyzer is 10.4 nm. For this size, when compared, it is expected that the transport rate is close to the nuclear uptake rate of similar-sized dextran molecules [7, 8, 6, 37]. However, the unexpectedly high value of nuclear import rates and transport behaviour gives us a new perspective on the nuclear transport of charged nanoparticles and the interaction of the charged particle with FG Nups.

## 2 Materials and methods

### 2.1 Materials

The chemicals and reagents used are of analytical grade. The Dulbecco’s Modified Eagle Medium (DMEM), Fetal Bovine Serum (FBS), Trypsin-EDTA, Dulbecco’s Phosphate buffer saline (PBS), Antibiotic antimycotic solution, N-(2-Hydroxy ethyl)-piperazine ethane sulfonic acid (HEPES), Potassium Acetate (KAc), Sodium Acetate (NaAc), Magnesium Acetate (MgAc), Ethylene glycol-bis(2-aminoethyl ether)-N,N,N’,N’-tetra acetic acid (EGTA), 1,4 Dithiothreitol (DTT), Protease Inhibitor, Digitonin are purchased from Himedia, India. RRL is obtained from Promega, U.S.A. The transport buffer contained 20 mM HEPES, 110 mM KAc, 5 mM NaAc, 2 mM MgAc, and 0.5 mM EGTA, pH 7.3. The complete transport buffer is made by adding 2mM DTT and 1 μg/ml of Protease inhibitor (aprotinin, leupeptin, and pepstatin) to the transport buffer, maintaining the same pH value. Digitonin stock solution is prepared by mixing 40 mg of digitonin powder in 1 ml of dimethyl sulfoxide. The stock solution is further diluted using the complete transport buffer and 40 μg/ml digitonin solution is used for permeabilization.

For the experiments with transport buffer containing RRL, the import solution is made by mixing 12.5 μl dialyzed RRL, 10 μl complete transport buffer, and 2.5 μl of GQDs solution. Whereas experiments carried out with transport buffer not containing RRL, the import solution comprised of 2.5 μl GQDs and 22.5 μl of complete transport buffer (GQDs with a stock concentration of 2 mg/ml).

### 2.2 Cell Culture

The HeLa cell line used in the current study is purchased from the Cell Repository division of the National Centre for Cell Science (NCCS), India. The cells are cultured in DMEM supplemented with 10% FBS, 50U/mL penicillin, and 0.05mg/mL streptomycin and incubated at 37 °C and 5% CO_2_. For live cell imaging, the cells are grown in an imaging chamber (confocal dish) with around 20,000 cells in 100 μl complete media and incubated for 12 hours prior to the experiment. The imaging chambers used are made in-house. For preparing the imaging chamber, a 6 mm hole is drilled in a 35 mm petri dish, and then a coverslip is fixed to the bottom of the petri dish using melted parafilm.

### 2.3 Experimental methods

To begin with, the imaging chamber containing cells is placed on the microscope stage of a confocal fluorescence microscope (FV 3000, Olympus Corporation) equipped with a 60X oil immersion objective and observed under the bright field. Transport buffer is used for washing the cells three times to remove any cell debris or unattached cells. After washing, cells are treated with digitonin for 5 minutes to permeabilize the cell membrane. After permeabilization, the cells are washed thrice with 30 μL of complete transport buffer. The nuclei can be seen clearly and distinctly after the permeabilization. The import mixture is prepared as per the description given in the materials section. The confocal laser scanning microscope is started in the timelapse mode of the 405 nm channel just before the addition of the import mixture. The transport buffer is carefully removed from the well, and the import mixture is added without disturbing the position of the well. The time-lapse confocal images are acquired for 5 minutes at the rate of 1.6 seconds per frame. It captures the fluorescence intensity from a confined area (central cross-section of the HeLa nuclei) as a function of time. Once the nuclear import achieves saturation, time-lapse export studies are performed. For this, the excess import mixture is wicked off completely, and 60 μl complete transport buffer is added after starting the time-lapse imaging. At this stage, the concentration of GQDs is high inside the nucleus compared to the outside. As a result, the GQDs start diffusing from the nucleus to the outside volume. Time-lapse confocal imaging is carried out for 5 minutes to capture this diffusion. To verify the retention properties of GQDs inside the nucleus, multiple export studies are carried out. For this, after completion of the first export study as described above, the transport buffer solution is removed completely, and again fresh complete transport buffer is added to the imaging chamber. This assures that the concentration of GQDs outside of the nucleus is zero at the beginning of each export study. Time-lapse confocal imaging is carried out again in this manner to see whether there is any further diffusion of GQDs from the nucleus. This export protocol is repeated three times.

To verify that the nucleus is intact during imaging, a control experiment is carried out at the end of each export experiment. TRITC-labeled dextran molecules of molecular weight 70 kDa having a size much larger than the passive diffusion limit are used for this. The experiment is carried out by adding the import mixture containing 70 kDa dextran to the imaging chamber and monitoring the fluorescence increase inside the nucleus. An intact nucleus will not show any increase in fluorescence inside the nucleus with time and will appear dark.

We have used Fiji (Image J), an open-source software for image analysis. We have analyzed the raw images by measuring the fluorescence intensity inside the nucleus, excluding the nuclear envelope. Normalized fluorescence is calculated by taking the ratio of fluorescence intensity inside the nucleus to that of outside.

## 3 Results

### 3.1 Nuclear import studies

Studies on the nuclear import and export of graphene quantum dots (GQDs) are carried out in digitonin permeabilized HeLa cells using time-lapse confocal fluorescence microscopy. The autofluorescence property of the GQDs is use of to detect them using the 405 nm excitation wavelength of the confocal microscope. Confocal imaging of the central cross-section of the nuclei is started before the addition of the complete transport buffer containing the GQDs. Fig. 1 shows representative images of the central cross-section of the nuclei at different times after the addition of the import mixture containing GQDs as well as RRL. The figure shows that the GQDs rapidly enter the nucleus and distribute themselves inside the nucleus. It may be noted that the diffusion rate inside the nucleoplasm is much higher than the transport rate through the nuclear membrane. In this study, we are focusing on the transport rate through the nuclear pore complex. Also, it is observed that nuclear organelles like nucleoli and nuclear membranes are brighter than the other parts of the nucleoplasm. The nuclear membrane is excluded during the analysis of the kinetics of transport to the nucleus.

**Figure 1:**
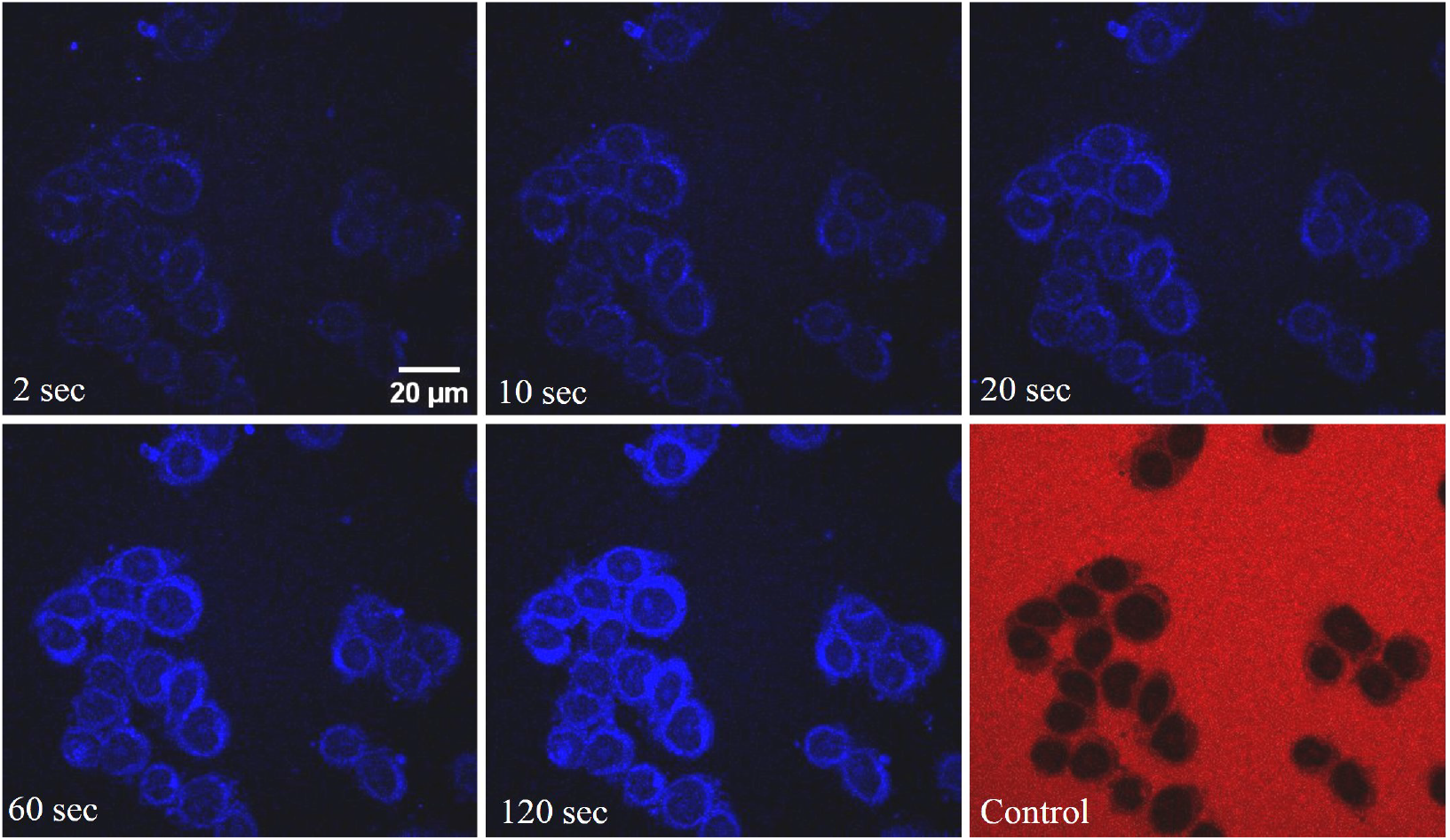
Representative time-lapse confocal images depicting the nuclear uptake of GQDs in HeLa cell nuclei. The frames show the image of the central cross-section of the nuclei after the addition of the import mixture containing RRL, at times indicated in the image. The last frame shows the results of a control experiment where the transport of 70 kDa TRITC dextran to the nucleus is monitored. The scale bar is 20 μm.

The fast nuclear translocation of GQDs into the nucleus for an import mixture containing RRL may, at first glance, indicate some form of active nuclear transport involving the proteins present in the RRL. To verify whether the proteins present in RRL play a role in nuclear uptake, the nuclear import experiment is performed using an import mixture not containing RRL. Fig. 2 shows a representative time-lapse image of the nuclear import carried out using an import mixture not containing RRL. The images shown in Fig. 2 indicate that the nuclear uptake behavior is similar to the one with the import mixture having RRL. The translocation rate in both cases is nearly the same (Table 1).

**Table 1:** The rate constant of transport of GQDs in HeLa cell nuclei determined from the nuclear import and export experiments.

| Nuclei type | Rate constant $k_1$ | Rate constant $k_2$ |
| --- | --- | --- |
| HeLa nuclei - import studies with RRL | $0.31 \pm 0.14$ | $0.008 \pm 0.003$ |
| HeLa nuclei - import studies without RRL | $0.28 \pm 0.14$ | $0.008 \pm 0.004$ |
| HeLa nuclei - export studies | $0.30 \pm 0.14$ | $0.010 \pm 0.004$ |

**Figure 2:**
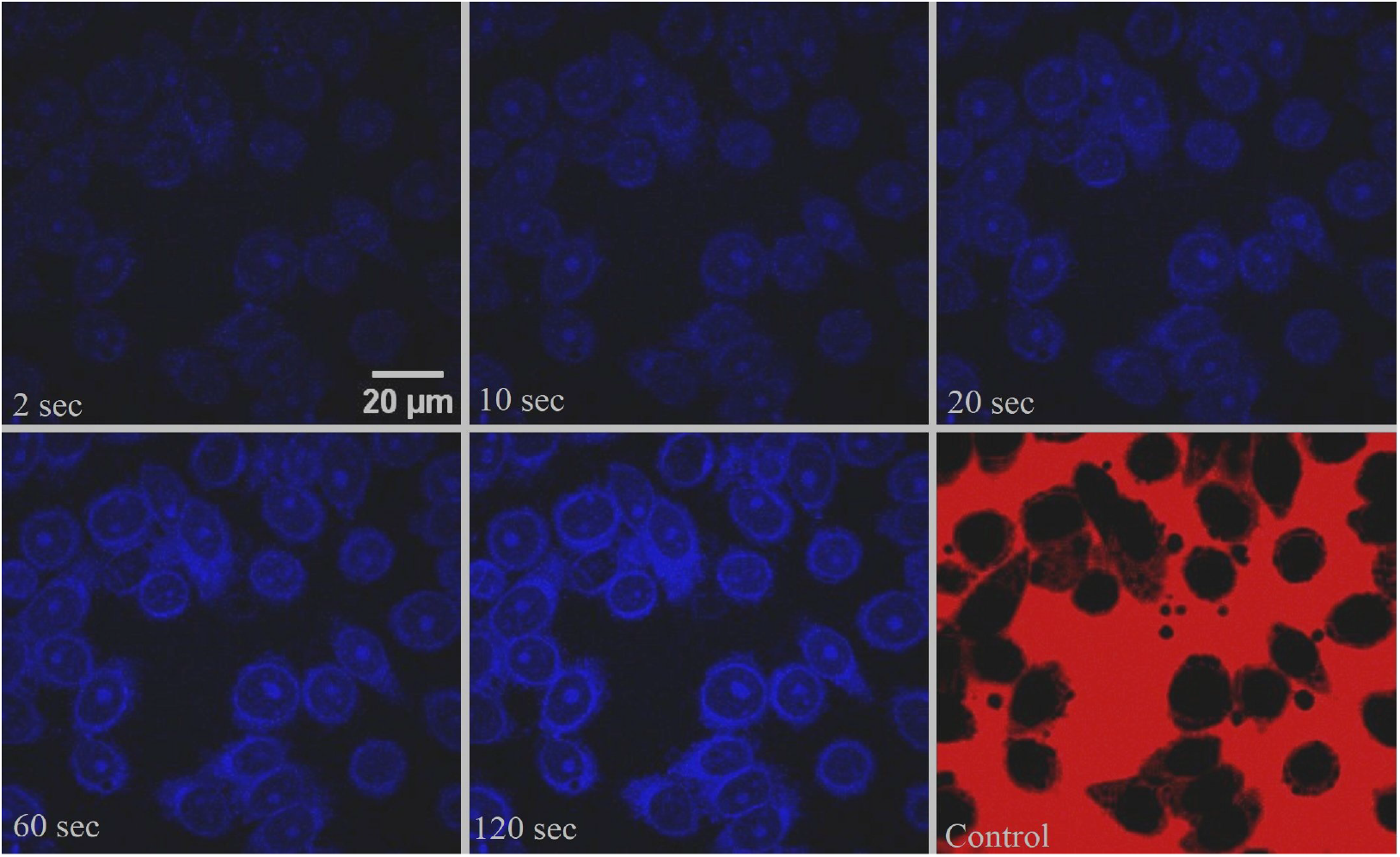
Time-lapse confocal images depicting the nuclear uptake of GQDs in HeLa cell system not containing RRL. The last frame shows the results of a control experiment carried out with 70 kDa TRITC dextran. Scale bar is 20 μm.

The HeLa cells employed in our studies are derived from the cervical cancer cell line. Tumors have a unique microenvironment characterized by hypoxia, inflammation, and angiogenesis that can impact the effectiveness of drug delivery using nanoparticles. The abnormal architecture and enhanced permeability of the tumor tissue can lead to increased retention and accumulation of nanoparticles, potentially improving the therapeutic efficacy of cancer treatments [38, 39].

It is possible that the enhanced import rate of nanoparticles in cancer cell nuclei may be due to the specific behavior of cancer cells towards nanoparticles. To check whether the uptake behaviour is due to the enhanced permeability and retention effect of cancer cells, experiments are conducted in HEK 293 nuclei. We find that the nuclear uptake behaviour in HEK 293 cell nuclei is similar to that of HeLa nuclei (Fig. 3). Import studies on HEK 293 nuclei establish that the fast translocation of GQDs to the HeLa cell nuclei is not due to the specific characteristics of cancer cells. The interaction of GQDs with NPCs plays an important role in nuclear uptake.

**Figure 3:**
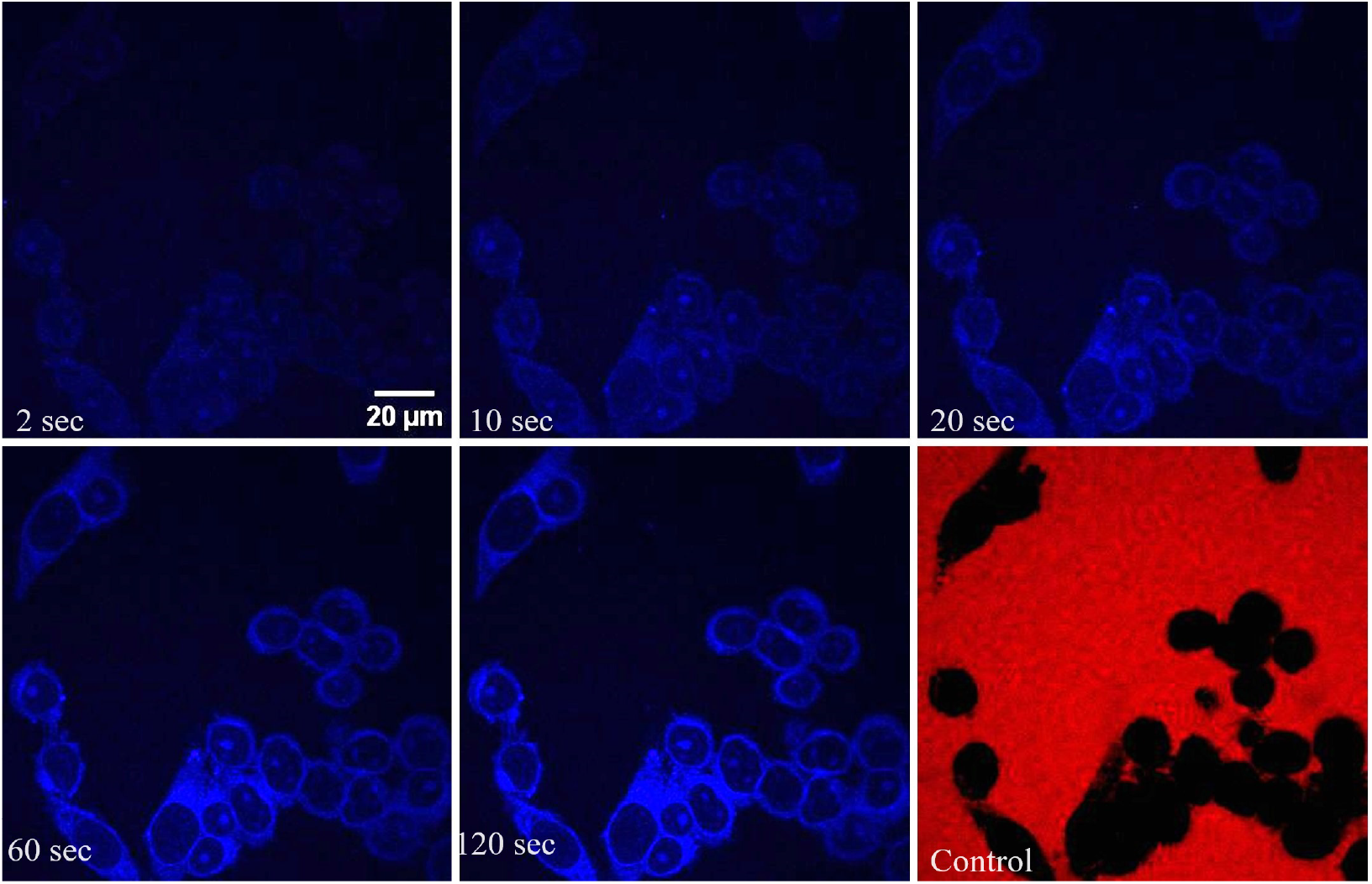
Nuclear uptake of GQDs in HEK 293 cell nuclei. The last frame shows the results of a control experiment carried out with 70 kDa TRITC dextran. Scale bar is 20 μm.

### 3.2 Modelling of the transport

The size of GQDs used for the study is within the passive diffusion limit, and NLS or importin *β* is not attached to GQDs. It is expected that the GQDs are transported to the nucleus via a passive diffusion mechanism, obeying a first-order kinetic equation of the form

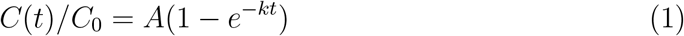

Here *C*(*t*) is the concentration of GQDs inside the nucleus, and *C*_0_ is the concentration outside of the nucleus. *k* is the rate constant of transport. *A* is a constant that is equal to one in the case of passive diffusion. This passive diffusion model implies that the rate of transport from the cytoplasm to the nucleus is equal to the rate of transport from the nucleus to the cytoplasm. To analyze the kinetics of the transport, we determine the normalized fluorescence intensity inside the nucleus by taking the ratio of the average fluorescence intensity inside the nucleoplasm to the average fluorescence intensity outside the nuclei. Since the fluorescence intensity is proportional to the concentration of fluorescing molecules, the normalized fluorescence intensity is equal to the ratio *C*(*t*)*/C*_0_.

The concentration of GQDs used in the import mixture is 0.2 mg/ml. Since the volume outside of the nucleus in the imaging chamber is very large, it is assumed that the outside concentration remains constant. Thus,

GQDs concentration inside nucleus = 0.2 mg/ml *×* Normalized fluorescence intensity

In Fig.4, we plot the normalized fluorescence intensity inside the nucleus as a function of time. It is observed that the data does not fit well with the single exponential behavior described by equation (1). The deviation of import data from equation (1) may indicate the presence of more than one independent transport pathway. If two independent transport pathways exist, the normalized concentration inside the nucleus as a function of time may be expressed as

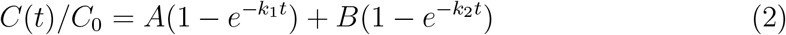

where C(t) is the concentration of graphene quantum dots inside the nucleus as a function of time, and *C*_0_ is the concentration of graphene quantum dots outside the nucleus. A is the normalized nuclear fluorescence of the initial fast component having rate *k*_1_ and B is the normalized nuclear fluorescence intensity corresponding to the slow component having rate *k*_2_ at the nuclear import saturation state. In Fig. 4 we show the fit of the experimental nuclear transport data (dots) to equation (2) (continuous line). The excellent agreement between the data and the fit indicates that the nuclear transport of GQDs follows a double exponential curve, and supports the assumption that two distinct mechanisms with different rate constants are involved in the transport. The experiment and analysis are repeated on a large number of nuclei, and histograms of the translocation rates, *k*_1_ and *k*_2_, of GQDs are plotted to determine the average rate constants (Fig. 5). The analysis reveals two distinct rate constants of transport, *k*_1_ = 0.31 *±*0.14 and *k*_2_ = 0.008 *±*0.003 in the case of GQDs transport in the presence of RRL. The rates in the absence of RRL in the import mixture are also the same, *k*_1_ = 0.28 *±*0.14 and *k*_2_ = 0.008 *±*0.004. The very similar values of rate constants in both these cases indicate that RRL does not play a significant role in transport. It may also be noted here that the rate constant *k*_2_ = 0.008 is of the order of the passive diffusion rate constant of dextran molecules of similar size while *k*_1_ is much higher [8, 37]. The import studies in HEK 293 nuclei show similar behaviour and rate constants.

**Figure 4:**
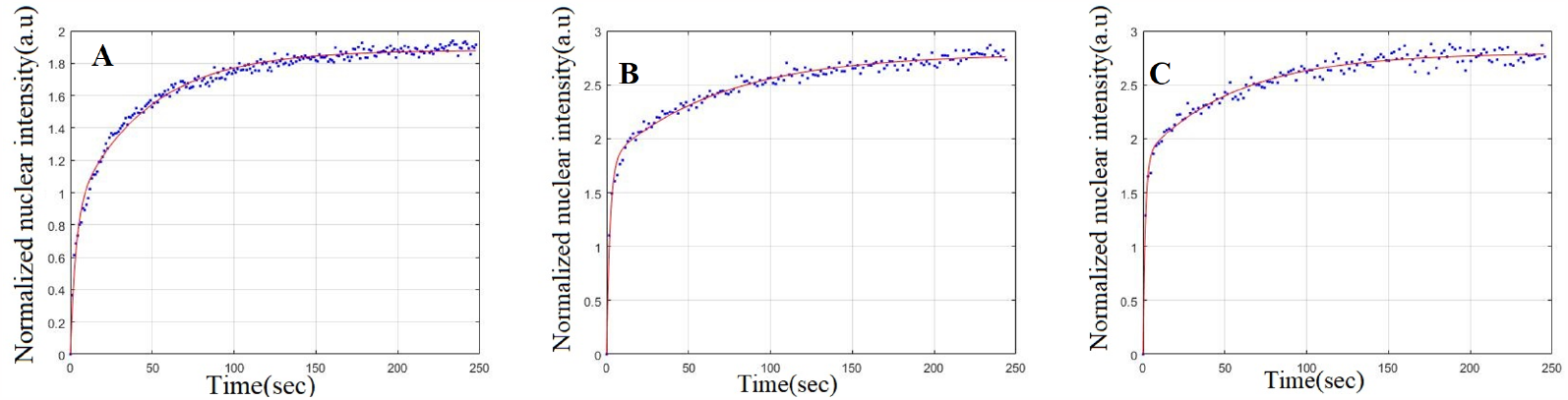
Normalized fluorescence intensity inside the nucleus as a function of time depicting the nuclear entry of GQDs in the case of (A) HeLa cell nuclei in the presence of RRL, (B) HeLa cell nuclei in the absence of RRL, and (C) HEK 293 cell nuclei in the absence RRL. The dots are experimental points, and the continuous line is fit to equation 2.

**Figure 5:**
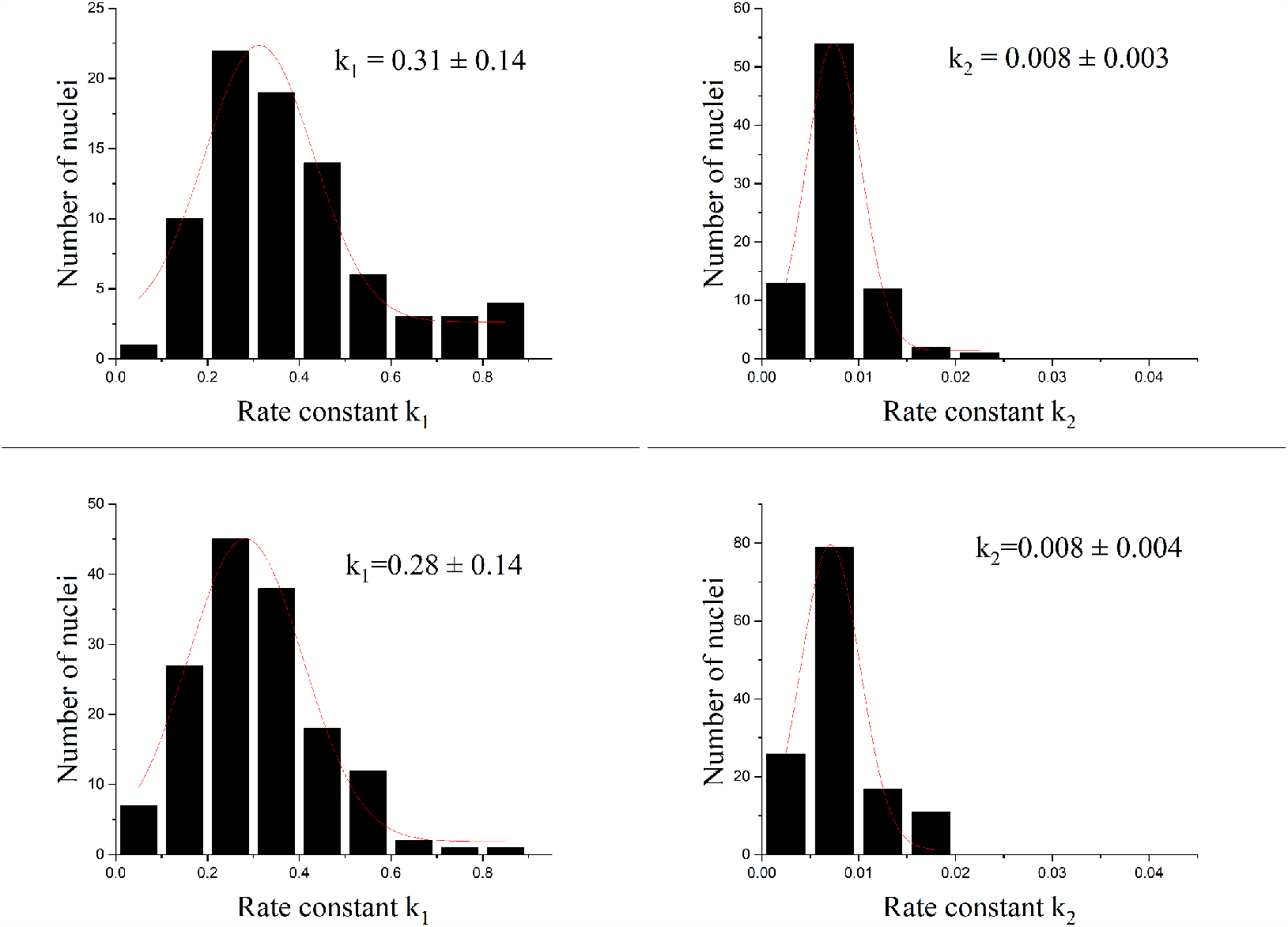
Histogram of the nuclear import rate constants *k*_1_ and *k*_2_ in HeLa nuclei A) with RRL and B) without RRL, respectively.

### 3.3 Maximum uptake intensity

In passive diffusion of biomolecules, the maximum uptake ratio (ratio of inside to outside fluorescence intensity) is expected to be 1. In these experiments it is observed that for GQDs, the ratio reaches the value of 3 (Fig. 4). This means GQDs concentration inside the nucleus is 3 times the concentration outside the nucleus. The concentration value of GQDs inside the nucleus saturates at around 0.5 mg/ml. An important question associated with this observation is whether the GQDs are getting attached to the nuclear components. Some weak interaction probably occurs between GQDs and nuclear components. From Fig. 1, 2, and 3, one can also observe that nucleoli are much brighter than the rest of the nucleus.

### 3.4 Nuclear Export studies

As discussed in the previous section, our data on nuclear import indicates that there are two distinct transport pathways for the nuclear uptake of GQDs. It is also observed that the concentration of GQDs inside the nucleus at the endpoint of the transport is much higher than that outside. It is important to know the reason for the above findings and the bi-directionality of interaction between GQDs and FG nups.To further verify and understand the above findings, we directly monitor the export of GQDs from the nucleus after the completion of the import assay. The export study is carried out by first removing the import mixture containing GQDs from the imaging chamber and adding fresh complete-transport buffer. Since the concentration of GQDs inside the nucleus is high at this point, the GQDs are exported from the nucleus to outside regions because of the concentration gradient. In Fig. 6, we show the time-lapse confocal images of the nuclei depicting the diffusion of GQDs from the nucleoplasm. It can be observed that the fluorescence intensity inside the nucleus decreases with time but does not reach zero. This indicates that GQDs do not diffuse out completely. Some amount of GQDs is retained in the nuclei after the export assay.

**Figure 6:**
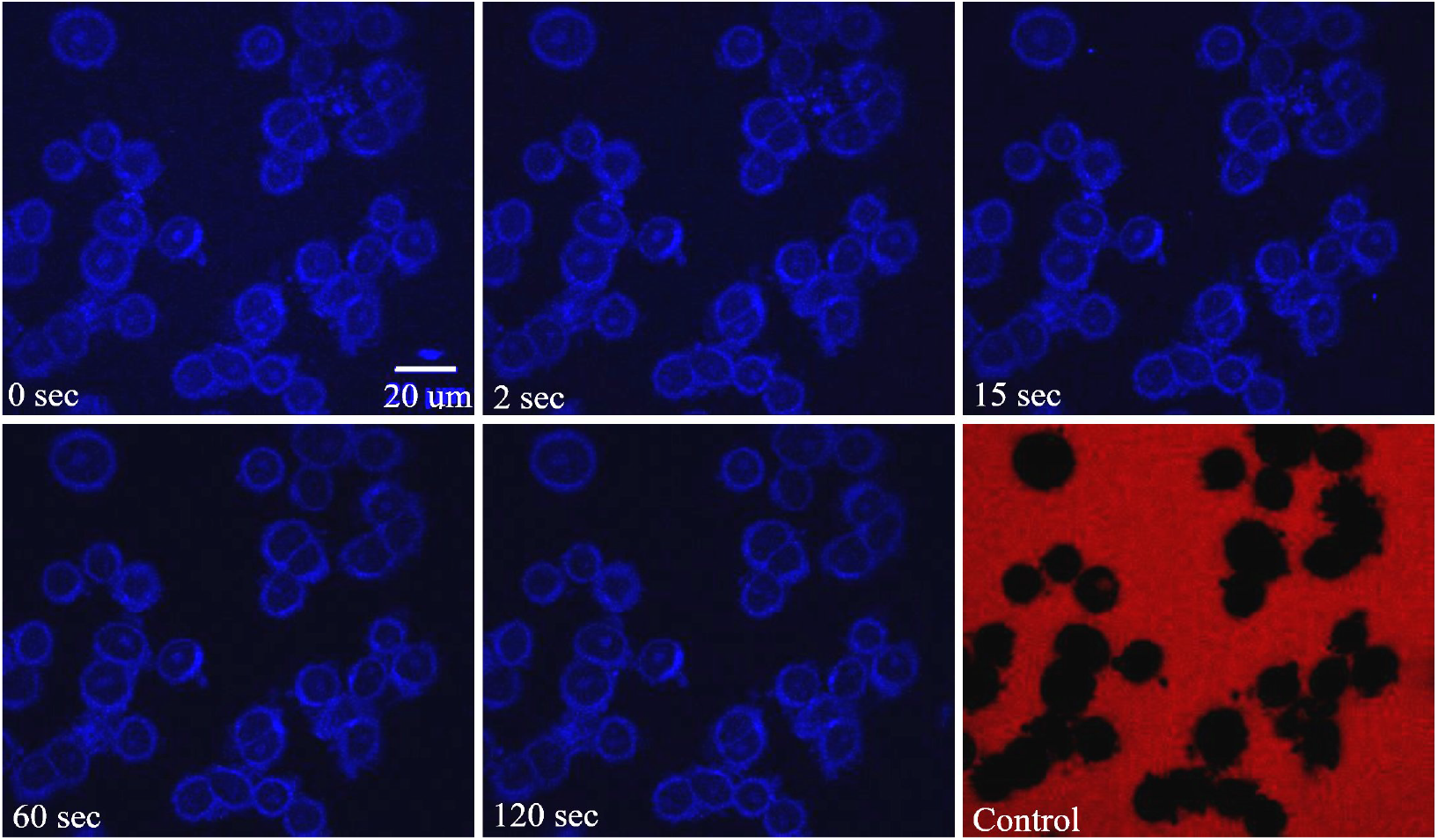
Time-lapse confocal images depicting the export of GQDs from HeLa cell nuclei. The frames show the images of the central cross-section of the nuclei at different times after the removal of the import mixture. The first image is taken after the import reached a steady state. The scale bar is 20 μm.

In the present case, diffusion takes place from the nucleus, where there is a high concentration of GQDs, to a much larger volume outside, where the concentration of GQDs can be approximated to zero. Hence the concentration *C*(*t*) inside the nucleus is expected to decrease exponentially,

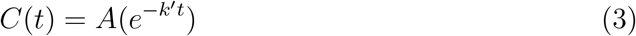

where *k*’ is the export rate constant. However, we find that fluorescence intensity inside the nucleus *C*(*t*) does not follow a single exponential curve. Hence we tried to fit the data to a double exponential decay given by the following equation

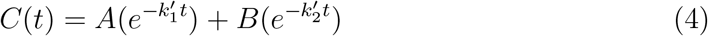

Here A and B are the maximum concentrations of graphene quantum dots inside the nucleus at the beginning of the export study corresponding to fast and slow export rate constants.

Fig. 7A shows that the experiment agrees well with the above model, implying that there are two different export pathways similar to what is observed in the nuclear import study. Fig. 7B and Fig. 7C depict the histogram of the export rate constants obtained by studying a large number of nuclei. The export rate constants obtained from the analysis are 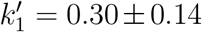 and 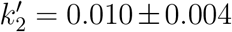. This indicates that the transport rate constants obtained from both the import and export studies are very similar. We have mentioned earlier that the higher import rate constant could be due to the interaction between the GQDs and FG Nups. This result leads us to one more finding that the same interaction occurs during the export process, and that the translocation of GQDs is bidirectional.

**Figure 7:**
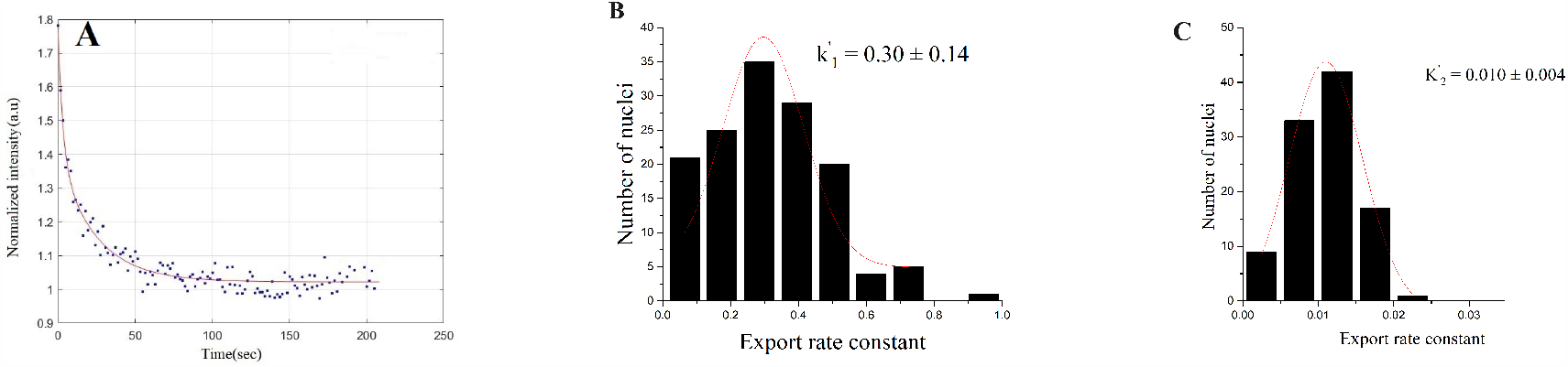
A) Normalized fluorescence intensity inside the nucleus is plotted as a function of time (dots) for nuclear export studies. The solid line is a fit to the double exponential decay curve given by equation 4. B) Histogram of the rate constant 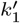 C) histogram of the rate constant 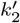.

### 3.5 Multiple export studies and retention

From export studies, it is found that the GQDs are not exported completely. To check if GQDs are getting drained off the nucleus after multiple washes, multiple export experiments are performed. For multiple export studies, we perform three consecutive export studies on the same set of nuclei.

Image A, B, and C in Figure 8 depict a consistent fluorescence intensity within the nucleus. These images represent the final frames of the first, second, and third export studies, respectively. The consistent fluorescence intensity indicates that there is no alteration in the concentration of GQDs, even after several attempts to export them. We plot a histogram of the retained GQDs concentration within the nucleus (Figure 8F). The images displayed in Fig. 8 affirm that GQDs remain retained within the nucleus, even following multiple washing procedures. The average concentration of GQDs retaining inside the nucleus after multiple washes is 0.3 ± 0.1 gm/ml. The GQDs are present across the nucleus but more prominent in the nucleoli. The possible reason for the retention could be the electrostatic interactions between GQDs and biomolecules present in the nucleus.

**Figure 8:**
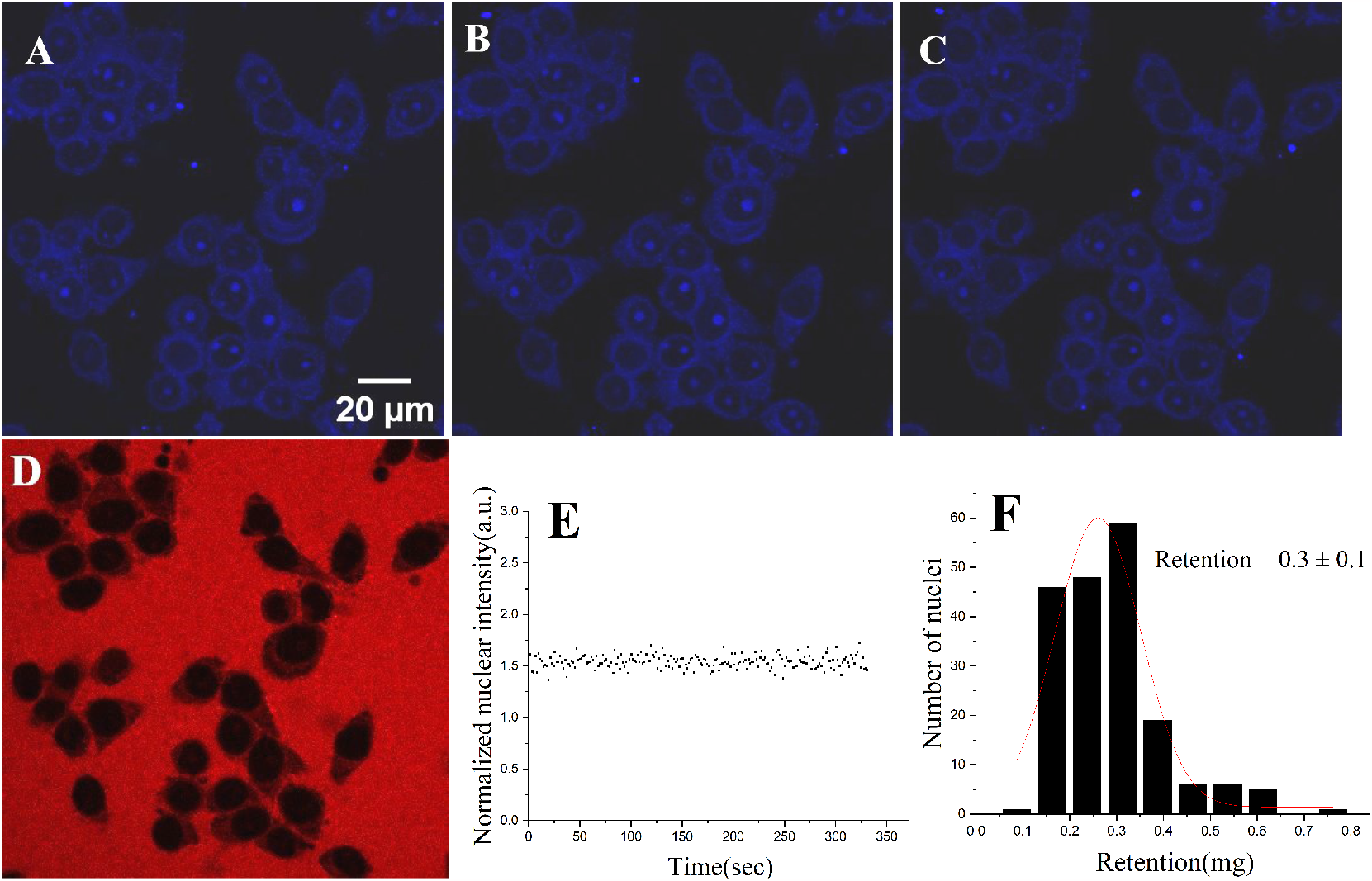
A) Image at the end of first export, B) Image at the end of second export, C) The last frame at the end of three export studies, D) Control experiment with 70 kDa dextran, scale bar is 20 μm, E) The graph of normalized nuclear intensity as a function of time, F) The histogram of retention in mg/ml.

## 4 Discussion

Quantum dots or nanoparticles, with their large surface-to-volume ratio, have a high potential to be a viable drug delivery vehicle. However, the transport of nanoparticles through cellular and nuclear membranes is not yet well studied and understood. There are few studies on the kinetics of nuclear transport of nanoparticles, and the translocation rates of nanoparticles through the nuclear membrane are seldom known. It is important to understand the interaction between nanoparticles and NPCs and to elucidate the role of surface charge and other parameters on the nuclear transport for the effective use of these particles in drug delivery and fundamental studies on the nuclear envelope. In this work, we study the transport of GQDs through the HeLa cell nuclear membrane and examine their transport characteristics. We used time-lapse confocal fluorescence microscopy to record the nuclear imports of GQDs in digitonin-permeabilized cells, and the rate constants of transport are determined using a simple diffusion-based model. The transport rates are further confirmed and verified via an independent experiment on nuclear export.

Our results demonstrate that the transport behavior of GQDs across the nuclear membrane differs significantly from that of standard probes such as FITC-dextran. The differences are noted in terms of faster nuclear translocation and retention of GQDs in the nucleus.

Two translocation rates, one fast and one slow, observed in the nuclear transport of GQDs lead to the interpretation that two phenomena are occurring simultaneously during the nuclear uptake. The fact that the rate constant of the slow component of transport is identical to that of passively diffusing molecules of similar size suggests that this is due to passive diffusion (diffusing through FG Nups without any interaction) [8, 37]. On the contrary, the dominating fast component could be attributed to transport mechanisms influenced by the surface properties of nanoparticles and their interactions with FG Nups.

The mechanism of active nuclear transport involves several complex processes. The NLS containing cargo is attached with importins, and the transport through the NPCs occurs via specific interaction of importins with FG repeats. Importins contain negatively charged regions that interact with positively charged regions of FG Nups. Additionally, importins contain hydrophobic regions that can interact with FG repeats present on Nups lining the NPCs. These FG Nups are hydrophobic in nature and contain small positive charges on them [5, 12, 19, 13, 14]. Therefore, the interaction between importins and FG Nups can be based on a hydrophobic interaction, electrostatic interaction, or a combination of both. The presence of different interactions is nonadditive; which is why the independent study of charged hydrophilic biomolecule and neutral hydrophobic biomolecule is not enough to describe the path of charged hydrophobic biomolecule [21].

The GQDs used in the study do not have NLS attached to them, and the possibility of them interacting with importins present in the RRL via a conventional pathway is rare. Thus we may assume that active transport in the conventional sense does not occur in the case of GQDs. The observation that the rate constants are not much influenced by the presence of RRL in the import mixture also supports this hypothesis. As discussed earlier, the FG Nups in the NPCs that act as a barrier for transport have a positive charge, and all transport factors are negatively charged hydrophobic biomolecules [22, 20, 21, 16]. The presence of charge on the FG Nups and transporting molecules could have a bearing on the transport rates. The electrostatic interaction between the transport factors and FG Nups results in a gain of several units of *k*_*B*_*T* of energy, leading to a lowering of the translocation energy barrier and increased translocation rate [21, 18]. A negative molecular surface charge increases the nuclear pore binding probability and reduces the transport time for passive diffusion through the NPCs. The green fluorescent proteins with a negative charge have shorter translocation time as compared to the positively charged green fluorescent proteins of similar size. It is also noted that the charge cannot give any relaxation to the passive diffusion cut-off size limit [22, 20]. Our findings that the nuclear uptake of GQDs is not influenced by the contents of RRL would mean that the interaction of GQDs with the nucleus is electrostatic. The nuclear uptake studies carried out in HEK 293 cells indicate that the interaction of GQDs with the nucleus is independent of the cell line and is not a special behavior specific to cancer cells. Thus the electrostatic interactions may play a crucial role in the efficient translocation of GQDs into the nucleus.

Additionally, export studies confirm the bidirectional nature of transport. The results from the export experiments revealed that there are two export rate constants, and both export rates are comparable to the corresponding rate constant obtained from the import studies. This observation suggests that the electrostatic interaction between GQDs and FG Nups occurs during transport in both directions. It is noteworthy that this interaction is not influenced by transport factors. We are proposing signal-less and electrostatic interaction-based translocation within the passive limit. The fact that both the import and export rates are similar confirms the occurrence of signal-less but interaction-based translocation.

Our findings suggest that the two transport rates observed are due to two different mechanisms, one due to electrostatic interaction and the other due to non-interactive diffusion within the passive limit, and these are largely uncoupled. The bidirectional diffusion suggests that interactive and non-interactive diffusion are similar in both nucleocytoplasmic directions. The higher concentration of GQDs inside the nucleus than outside, as seen in Fig.4, can be attributed to the electrostatic interaction between the negatively charged GQDs and the positively charged biomolecules or any hydrophilic linker on the biomolecules present in the nucleus. The GQDs tend to form the *π* − *π* stacking or electrostatic bonding with biomolecules present in the nucleus. The nucleus comprises a complex network of biomolecules with varying charges, and these differences in charge create opportunities for the GQDs to bind and remain trapped within the nucleus. Additionally, the liquid-liquid phase separation within the nucleus may also result in small gaps or voids where the GQDs may be trapped, resulting in a higher concentration of GQDs within the nucleus. Many groups found that the uptake and retention of the cargo molecule within the nucleus are dependent on the interaction between the cargo’s charge and surface properties, composition, and charges of the biomolecules present in the nucleus [27]. Considering the surface properties of nanoparticles, it is plausible that interaction between GQDs and nuclear entities leads to better retention inside the nucleus [34].

The multiple export studies confirm the retention inside the nucleus. The fact that the GQDs are retained in the nucleoli region highlights the significance of the surface properties and electrostatic interactions between the GQDs and the biomolecules present in nucleoli. The GQDs may tend to attach with DNA through *π* − *π* stacking, and electrostatic interaction. The GQDs may interact with different biomolecules through hydrogen bonding and amide bond formation because there is the presence of carboxyl group[34, 40, 41]. The retention in the nucleoli needs further investigation to understand the precise mechanisms behind this retention and its potential applications in the field of biomedicine.

Our study shows that negatively charged nanoparticles can be an important carrier to enhance the nuclear uptake rate of biomolecules. The fast transport rates exhibited by GQDs show their potential utility as drug carriers, and the retention properties ensure the passing of the drug to daughter nuclei.

## 5 Conclusion

In the present work, we use time-lapse confocal fluorescence microscopy to study the kinetics of the transport of negatively charged GQDs through the nuclear membrane. The studies reveal a rapid nuclear uptake of GQDs, pointing to the potential application of GQds as a drug carrier. Results suggest that the nuclear uptake of GQDs is not influenced by the contents of RRL and that the high nuclear transport rates observed could be due to electrostatic interaction. The nuclear uptake studies carried out in HeLa cells and HEK 293 cells show that the interaction of GQDs with the nucleus is independent of the cell line and is not a special behavior specific to cancer cells.

The export studies confirm the bidirectional nature of transport. The results from the export experiments reveal that there are two export rate pathways. The analysis of import and export data reveals that the rate constants of import and export are comparable and thus suggests a bidirectional transport with similar kind of electrostatic interaction between GQDs and FG Nups occurring during transport in both directions.

The multiple export studies confirm good retention of GQDs inside the nucleus. The retention can make the drug remain in the cells and pass to every daughter cell from the mother cell.

## Author contributions

G.G., G.K.V., and P.N. conceived and designed the study. G.G. and V.K. maintained the cell cultures and conducted imaging experiments. G.G. analyzed and interpreted the fluorescence data along with G.K.V. and P.N. G.G., V.K., G.K.V., and P.N. wrote the manuscript.

## Acknowledgments

We would like to thank SD Hiremath and A Chatterjee, Chemistry department, BITS Goa, for providing the graphene quantum dots. We are grateful to Toby Joseph, Department of Physics, BITS Pilani K. K. Birla Goa campus, and M.M. Bijeesh, NTU Singapore, for their help in modeling and simulation of the nuclear transport. We are also thankful to Shakhi P.K. for introducing us to the field and initial work. Gorav and Vrushali are thankful to BITS Pilani for the research fellowship. Dr Geetha K Varier would like to thank the Department of Science and Technology for financial support under the DST-WOS-A scheme (WOS-A/PM-32/2018). The authors acknowledge the Central Sophisticated Instrumentation Facility (CSIF), BITS Pilani, Goa Campus for confocal laser scanning microscopy, SEM, DLS, and Zeta potential analysis; and the National Institute of Technology, Surathkal, Department of Metallurgical and Materials Engineering for TEM studies. Graphical abstract is partially made in biorender(Created with BioRender.com).

## References

[1] T. D. Allen, J. Cronshaw, S. Bagley, E. Kiseleva, M. W. Goldberg, The nuclear pore complex: mediator of translocation between nucleus and cytoplasm, Journal of cell science 113 (10) (2000) 1651–1659.

[2] D. Stoffler, B. Fahrenkrog, U. Aebi, The nuclear pore complex: from molecular architecture to functional dynamics, Curr Opin Cell Biol 11 (3) (1999) 391–401.

[3] S. A. Adam., The nuclear pore complex, Genome biology 2 (9) (2001) 1–6.

[4] A. M. Brownawell, J. M. Holaska, I. G. Macara, B. M. Paschal, The use of permeabilized cell systems to study nuclear transport, Methods Mol Biol 189 (2002) 209–229.

[5] I. G. Macara, Transport into and out of the nucleus, Microbiology and molecular biology reviews 65 (4) (2001) 570–594.

[6] O. Keminer, R. Peters, Permeability of single nuclear pores, Biophysical journal 77 (1) (1999) 217–228.

[7] D. Mohr, S. Frey, T. Fischer, T. Güttler, D. Görlich, Characterisation of the passive permeability barrier of nuclear pore complexes, The EMBO journal 28 (17) (2009) 2541–2553.

[8] A. Samudram, B. M. Mangalassery, M. Kowshik, N. Patincharath, G. K. Varier, Passive permeability and effective pore size of hela cell nuclear membranes, Cell Biology International 40 (9) (2016) 991–998.

[9] B. L. Timney, B. Raveh, R. Mironska, J. M. Trivedi, S. J. Kim, D. Russel, S. R. Wente, A. Sali, M. P. Rout, Simple rules for passive diffusion through the nuclear pore complex, Journal of Cell Biology 215 (1) (2016) 57–76.

[10] E. Dultz, J. Ellenberg, Live imaging of single nuclear pores reveals unique assembly kinetics and mechanism in interphase, Journal of Cell Biology 191 (1) (2010) 15–22.

[11] G. G. Maul, H. M. Maul, J. E. Scogna, M. W. Lieberman, G. S. Stein, B. Y.-L. Hsu, T. W. Borun, Time sequence of nuclear pore formation in phytohemagglutinin-stimulated lymphocytes and in hela cells during the cell cycle, The Journal of cell biology 55 (2) (1972) 433–447.

[12] K. Ribbeck, D. Görlich, Kinetic analysis of translocation through nuclear pore complexes, The EMBO journal 20 (6) (2001) 1320–1330.

[13] G. Kabachinski, T. U. Schwartz, The nuclear pore complex–structure and function at a glance, Journal of cell science 128 (3) (2015) 423–429.

[14] M. Beck, E. Hurt, The nuclear pore complex: understanding its function through structural insight, Nature Reviews Molecular cell biology 18 (2) (2017) 73–89.

[15] L. J. Terry, S. R. Wente, Flexible gates: dynamic topologies and functions for fg nucleoporins in nucleocytoplasmic transport, Eukaryotic cell 8 (12) (2009) 1814–1827.

[16] J. Yamada, J. L. Phillips, S. Patel, G. Goldfien, A. Calestagne-Morelli, H. Huang, R. Reza, J. Acheson, V. V. Krishnan, S. Newsam, et al., A bimodal distribution of two distinct categories of intrinsically disordered structures with separate functions in fg nucleoporins, Molecular & Cellular Proteomics 9 (10) (2010) 2205–2224.

[17] R. Hayama, S. Sparks, L. M. Hecht, K. Dutta, J. M. Karp, C. M. Cabana, M. P. Rout, D. Cowburn, Thermodynamic characterization of the multivalent interactions underlying rapid and selective translocation through the nuclear pore complex, Journal of Biological Chemistry 293 (12) (2018) 4555–4563.

[18] A. Ghavami, E. Van Der Giessen, P. R. Onck, Energetics of transport through the nuclear pore complex, PLoS One 11 (2) (2016) e0148876.

[19] G. Paci, J. Caria, E. A. Lemke, Cargo transport through the nuclear pore complex at a glance, Journal of cell science 134 (2) (2021) jcs247874.

[20] L. J. Colwell, M. P. Brenner, K. Ribbeck, Charge as a selection criterion for translocation through the nuclear pore complex, PLoS computational biology 6 (4) (2010) e1000747.

[21] M. Tagliazucchi, O. Peleg, M. Kröger, Y. Rabin, I. Szleifer, Effect of charge, hydrophobicity, and sequence of nucleoporins on the translocation of model particles through the nuclear pore complex, Proceedings of the National Academy of Sciences 110 (9) (2013) 3363–3368.

[22] A. Goryaynov, W. Yang, Role of molecular charge in nucleocytoplasmic transport, PloS one 9 (2) (2014) e88792.

[23] R. Mehvar, Dextrans for targeted and sustained delivery of therapeutic and imaging agents, Journal of controlled release 69 (1) (2000) 1–25.

[24] K. Nishida, K. Mihara, T. Takino, S. Nakane, Y. Takakura, M. Hashida, H. Sezaki, Hepatic disposition characteristics of electrically charged macromolecules in rat in vivo and in the perfused liver, Pharmaceutical research 8 (1991) 437–444.

[25] T. Yamaoka, M. Kuroda, Y. Tabata, Y. Ikada, Body distribution of dextran derivatives with electric charges after intravenous administration, International journal of pharmaceutics 113 (2) (1995) 149–157.

[26] J. Rouquette, C. Genoud, G. H. Vazquez-Nin, B. Kraus, T. Cremer, S. Fakan, Revealing the high-resolution three-dimensional network of chromatin and interchromatin space: a novel electron-microscopic approach to reconstructing nuclear architecture, Chromosome research 17 (2009) 801–810.

[27] T. Lebeaupin, R. Smith, S. Huet, The multiple effects of molecular crowding in the cell nucleus: from molecular dynamics to the regulation of nuclear architecture, Nuclear Architecture and Dynamics (2018) 209–232.

[28] C. Yang, J. Uertz, D. Yohan, B. Chithrani, Peptide modified gold nanoparticles for improved cellular uptake, nuclear transport, and intracellular retention, Nanoscale 6 (20) (2014) 12026–12033.

[29] W.-j. Jeong, J. Bu, L. J. Kubiatowicz, S. S. Chen, Y. Kim, S. Hong, Peptide– nanoparticle conjugates: a next generation of diagnostic and therapeutic platforms?, Nano Convergence 5 (1) (2018) 1–18.

[30] S. Silva, A. J. Almeida, N. Vale, Combination of cell-penetrating peptides with nanoparticles for therapeutic application: a review, Biomolecules 9 (1) (2019) 22.

[31] I. Gessner, I. Neundorf, Nanoparticles modified with cell-penetrating peptides: Conjugation mechanisms, physicochemical properties, and application in cancer diagnosis and therapy, International journal of molecular sciences 21 (7) (2020) 2536.

[32] U. Resch-Genger, M. Grabolle, S. Cavaliere-Jaricot, R. Nitschke, T. Nann, Quantum dots versus organic dyes as fluorescent labels, Nature methods 5 (9) (2008) 763–775.

[33] M. Nurunnabi, Z. Khatun, K. M. Huh, S. Y. Park, D. Y. Lee, K. J. Cho, Y.-k. Lee, In vivo biodistribution and toxicology of carboxylated graphene quantum dots, ACS nano 7 (8) (2013) 6858–6867.

[34] Z. Gu, S. Zhu, L. Yan, F. Zhao, Y. Zhao, Graphene-based smart platforms for combined cancer therapy, Advanced Materials 31 (9) (2019) 1800662.

[35] M. Karimi, S. Bahrami, S. B. Ravari, P. S. Zangabad, H. Mirshekari, M. Bozorgomid, S. Shahreza, M. Sori, M. R. Hamblin, Albumin nanostructures as advanced drug delivery systems, Expert opinion on drug delivery 13 (11) (2016) 1609–1623.

[36] R. Khandelia, S. Bhandari, U. N. Pan, S. S. Ghosh, A. Chattopadhyay, Gold nanocluster embedded albumin nanoparticles for two-photon imaging of cancer cells accompanying drug delivery, Small 11 (33) (2015) 4075–4081.

[37] M. Mangalassery, T. Joseph, N. Patincharath, G. Varier, et al., Size dependent steady-state saturation limit in biomolecular transport through nuclear membranes (2023).

[38] J. Fang, H. Nakamura, H. Maeda, The epr effect: unique features of tumor blood vessels for drug delivery, factors involved, and limitations and augmentation of the effect, Advanced drug delivery reviews 63 (3) (2011) 136–151.

[39] A. D. Wong, M. Ye, M. B. Ulmschneider, P. C. Searson, Quantitative analysis of the enhanced permeation and retention (epr) effect, PloS one 10 (5) (2015) e0123461.

[40] S. Rafiei, M. Dadmehr, M. Hosseini, H. A. Kermani, M. R. Ganjali, A fluorometric study on the effect of dna methylation on dna interaction with graphene quantum dots, Methods and applications in fluorescence 7 (2) (2019) 025001.

[41] L. Qi, T. Pan, L. Ou, Z. Ye, C. Yu, B. Bao, Z. Wu, D. Cao, L. Dai, Biocompatible nucleus-targeted graphene quantum dots for selective killing of cancer cells via dna damage, Communications Biology 4 (1) (2021) 214.

